# Clades of huge phage from across Earth’s ecosystems

**DOI:** 10.1101/572362

**Authors:** Basem Al-Shayeb, Rohan Sachdeva, Lin-Xing Chen, Fred Ward, Patrick Munk, Audra Devoto, Cindy J. Castelle, Matthew R. Olm, Keith Bouma-Gregson, Yuki Amano, Christine He, Raphaël Méheust, Brandon Brooks, Alex Thomas, Adi Lavy, Paula Matheus-Carnevali, Christine Sun, Daniela S. A. Goltsman, Mikayla A. Borton, Tara C. Nelson, Rose Kantor, Alexander L. Jaffe, Ray Keren, Ibrahim F. Farag, Shufei Lei, Kari Finstad, Ronald Amundson, Karthik Anantharaman, Jinglie Zhou, Alexander J. Probst, Mary E. Power, Susannah G. Tringe, Wen-Jun Li, Kelly Wrighton, Sue Harrison, Michael Morowitz, David A. Relman, Jennifer A Doudna, Anne-Catherine Lehours, Lesley Warren, Jamie H. D. Cate, Joanne M. Santini, Jillian F. Banfield

## Abstract

Phage typically have small genomes and depend on their bacterial hosts for replication. DNA sequenced from many diverse ecosystems revealed hundreds of huge phage genomes, between 200 kbp and 716 kbp in length. Thirty-four genomes were manually curated to completion, including the largest phage genomes yet reported. Expanded genetic repertoires include diverse and new CRISPR-Cas systems, tRNAs, tRNA synthetases, tRNA modification enzymes, translation initiation and elongation factors, and ribosomal proteins. Phage CRISPR-Cas systems have the capacity to silence host transcription factors and translational genes, potentially as part of a larger interaction network that intercepts translation to redirect biosynthesis to phage-encoded functions. In addition, some phage may repurpose bacterial CRISPR-Cas systems to eliminate competing phage. We phylogenetically define major clades of huge phage from human and other animal microbiomes, oceans, lakes, sediments, soils and the built environment. We conclude that their large gene inventories reflect a conserved biological strategy, observed over a broad bacterial host range and across Earth’s ecosystems.

---

Bacteriophage (phage), viruses of bacteria, are considered distinct from cellular life due to their inability to conduct most biological processes required for reproduction. They are agents of ecosystem change because they prey upon specific bacterial host populations, mediate lateral gene transfer, alter host metabolisms, and redistribute bacterially-derived compounds via cell lysis (Rascovan *et al.*, 2016; Breitbart *et al.*, 2018; Emerson *et al.*, 2018). They spread antibiotic resistance (Balcazar, 2014) and disperse pathogenicity factors that cause disease in humans and animals (Penadés *et al.*, 2015; Brown-Jaque *et al.*, 2018). Most knowledge about phage is based on laboratory-studied examples, the vast majority of which have genomes a few 10s of kbp in length. Widely used isolation-based methods select against large phage particles, and they can be excluded from phage concentrates obtained by passage through 100 nm or 200 nm filters (Yuan and Gao, 2017). In 2017, only 93 isolated phage with genomes of >200 kbp in length had been published (Yuan and Gao, 2017). This motivates cultivation-independent approaches to evaluate the existence and biology of phage with huge genomes. Only with this information on hand will it be possible to evaluate the roles that these phage play in the functioning and evolution of ecosystems.

Sequencing of whole community DNA can uncover phage-derived DNA fragments, yet large phage genomes can still escape detection due to genome fragmentation (Shkoporov and Hill, 2019). However, a new clade of human and animal-associated megaphage was recently described based on phage genomes manually curated to completion from metagenomic datasets (Devoto *et al.*, 2019). This finding motivated more comprehensive analysis of microbial communities to evaluate the prevalence, diversity and ecosystem distribution of phage with large genomes. Here, we present hundreds of phage sequences >200 kbp in length that we reconstructed from microbiome datasets generated from a wide variety of ecosystems. A graphical abstract provides an overview of our approach and main findings (**Figure S1**). We reconstructed the three largest complete genomes for phage known to date, ranging up to 642 kbp in length. The research expands our understanding of phage biodiversity and reveals the wide variety of ecosystems in which phage have genomes with sizes that rival those of small celled bacteria (Nakabachi *et al.*, 2006; Pérez-Brocal *et al.*, 2006; Castelle *et al.*, 2018). We postulate that these phage have evolved a distinct ‘life’ strategy that involves extensive interception and augmentation of host biology while they replicate their huge genomes.

## Ecosystem sampling

Metagenomic datasets were acquired from human fecal and oral samples, fecal samples from other animals, freshwater lakes and rivers, marine ecosystems, sediments, hot springs, soils, deep subsurface habitats and the built environment (**Table 1**). Genome sequences that were clearly not bacterial, archaeal, archaeal virus, eukaryotic or eukaryotic virus were classified as either phage or plasmid-like based on their gene inventories. *De novo* assembled fragments close to or >200 kbp in length were tested for circularization and a subset selected for manual verification and curation to completion (see Methods).

**Table 1:**
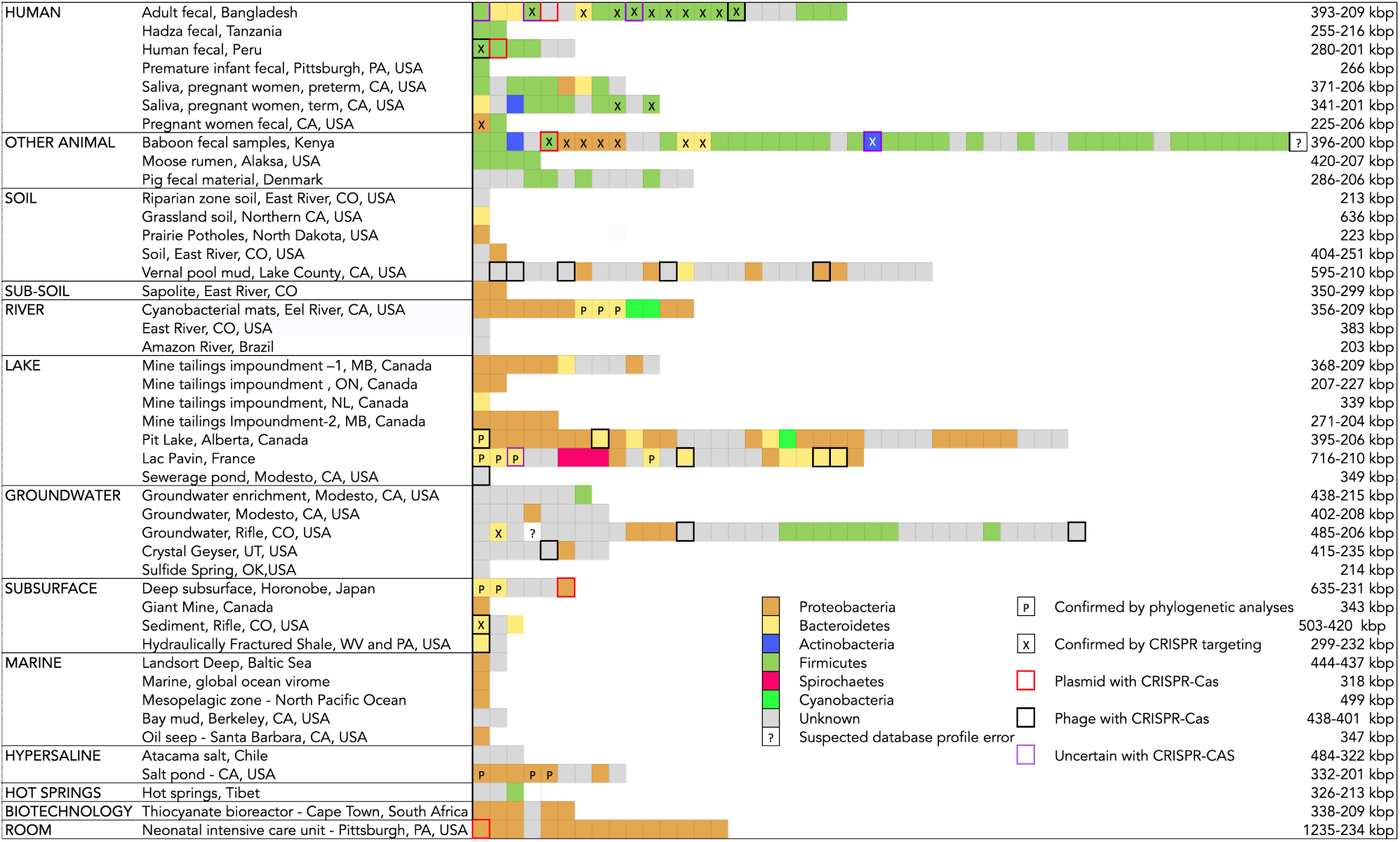
Ecosystems with phage and plasmid-like genomes >200 kb, grouped by sampling site type. Each box represents a genome, and boxes are horizontally arranged in order of decreasing genome size. Size range for each site type is listed to the right. Colors indicate putative host phylum based on genome taxonomic profile, with confirmation by CRISPR targeting (X) or information system gene phylogenetic analyses (T).

## Genome sizes and basic features

We reconstructed 353 phage, 5 plasmid-like, and 3 uncertain phage-plasmid sequences (**Table 1, S1**). We excluded additional sequences inferred to be plasmids (see Methods), retaining only those encoding CRISPR-Cas loci. We included 3 phage sequences of <200 kbp in length due to the presence of CRISPR-Cas loci. Consistent with classification as phage, we identified a wide variety of phage-relevant genes, including those involved in lysis and encoding structural proteins, and documented other expected phage genomic features (see Supplementary Information, SI). Some phage predicted proteins are large, up to 7,694 amino acids in length. Many of these were tentatively annotated as structural proteins. 180 phage sequences were circularized and 34 were manually curated to completion, in some cases by resolving complex repeat regions, revealing their encoded proteins (see Methods and **Table S1**). Approximately 30% of genomes show clear GC skew indicative of bi-directional replication, information that constrains their replication origin, and 30% indicative of unidirectional replication (SI and **Figure S2**) (Lobry, 1996).

Our three largest complete, manually curated and circularized phage genomes are 634, 636, and 642 kbp in length and represent the largest phage genomes reported to date. Previously, the largest circularized phage genome was 596 kbp in length (Paez-Espino *et al.*, 2016). The same study reported a circularized genome of 630 kbp in length, but this is an assembly artifact (SI and **Figure S3**). The problem of concatenated sequences was sufficiently prominent in IMG/VR (Paez-Espino *et al.*, 2017) that we did not include these data in further analyses. We used the complete and circularized genomes from our study and published phage genomes to depict a current view of the distribution of phage genome sizes (Methods). The median genome size for complete phage is ∼52 kbp (**Figure 1A**), similar to the average size of ∼54 kbp reported previously (Paez-Espino *et al.*, 2016). Thus, sequences reported here substantially expand the inventory of phage with unusually large genomes (**Figure 1B**).

**Figure 1:**
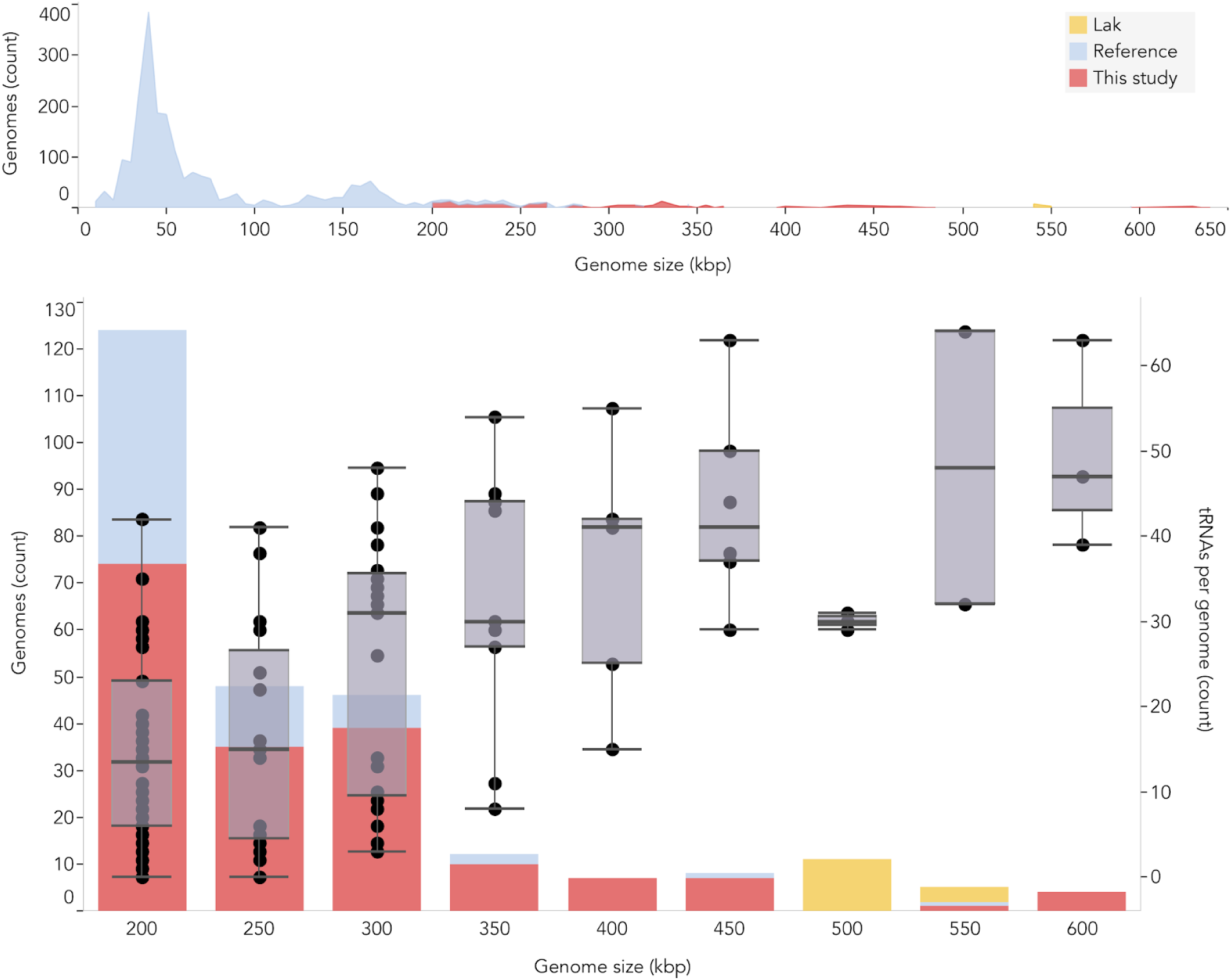
**A.** Size distribution of complete bacteriophage genomes from this study, Lak phage genomes reported recently from a subset of the same samples (Devoto *et al.*, 2019), and reference sources. Reference genomes were collected from all RefSeq r92 dsDNA genomes and nonartifactual assemblies >200 kb from (Paez-Espino *et al.*, 2016). **B.** Histogram of the genome size distribution of phage with genomes >200 kb from this study, Lak, and reference genomes. Box and whisker plots of tRNA counts per genome from this study and Lak phage as a function of genome size.

We identified and manually curated two related sequences of 712 and 716 kbp in length (**Table 1**), but the genomes could not be circularized due to a few kbp-long complex repeat regions at both genome ends. We anticipate that these genomes could be closed if the repeat regions could be rationalized. These were classified as phage sequences based on their overall genome content and the presence of terminase genes.

Some of our reported genomes have very low coding density (nine <78%, see SI), probably due to the use of a genetic code different from the standard code used for gene prediction (see Methods). This phenomenon has been rarely noted in phage, but was reported for Lak phage (Devoto *et al.*, 2019), and by (Ivanova *et al.*, 2014). In the current study, some genomes (mostly human/animal associated) appear to have reassigned the UAG (amber) stop codon to code for an amino acid (SI and **Figure S4**).

In only one case, we identified a sequence of >200 kbp that was classified as a prophage based on transition into flanking bacterial genome sequence. However, around half of the genomes were not circularized, so their potential integration as prophage cannot be ruled out. The presence of integrases in some genomes is suggestive of a lysogenic lifestyle under some conditions.

## Hosts, diversity, and distribution

An intriguing question relates to the evolutionary history of phage with huge genomes. Are they the result of recent genome expansion within clades of normal sized phage or is a large inventory of genes an established, persistent strategy? To investigate this, we constructed phylogenetic trees for large terminase subunit (**Figure 2**) and major capsid (**Figure S5A)** proteins using sequences in public databases as context (Methods). Many of the sequences from our large phage genomes cluster together with high bootstrap support, defining clades. Analysis of the genome size information for database sequences shows that the public sequences that fall into these clades are from phage with genomes of at least 120 kbp in length. The largest clade, referred to here as Mahaphage (Maha being Sanskrit for huge), includes all of our largest genomes as well as the 540 - 552 kbp Lak genomes from human and animal microbiomes (Devoto *et al.*, 2019). We identified nine other clusters of large phage, and refer to them using the words for “huge” in languages of some authors of this publication. We acknowledge that the detailed tree topologies for different genes and datasets vary somewhat, but the clustering is broadly supported by protein family and capsid analyses (**SI and Figure S5A**,**B**). The consistent grouping together of large phage into clades establishes that large genome size is a relatively stable trait (SI). Within each clade, phage were sampled from a wide variety of environmental types (**Figure 2**), indicating diversification of these large phage and their hosts across ecosystems. We also examined the environmental distribution of phage that are so closely related that their genomes can be aligned and found 19 cases where they occur in at least two distinct cohorts or habitat types (**Table S2**).

**Figure 2:**
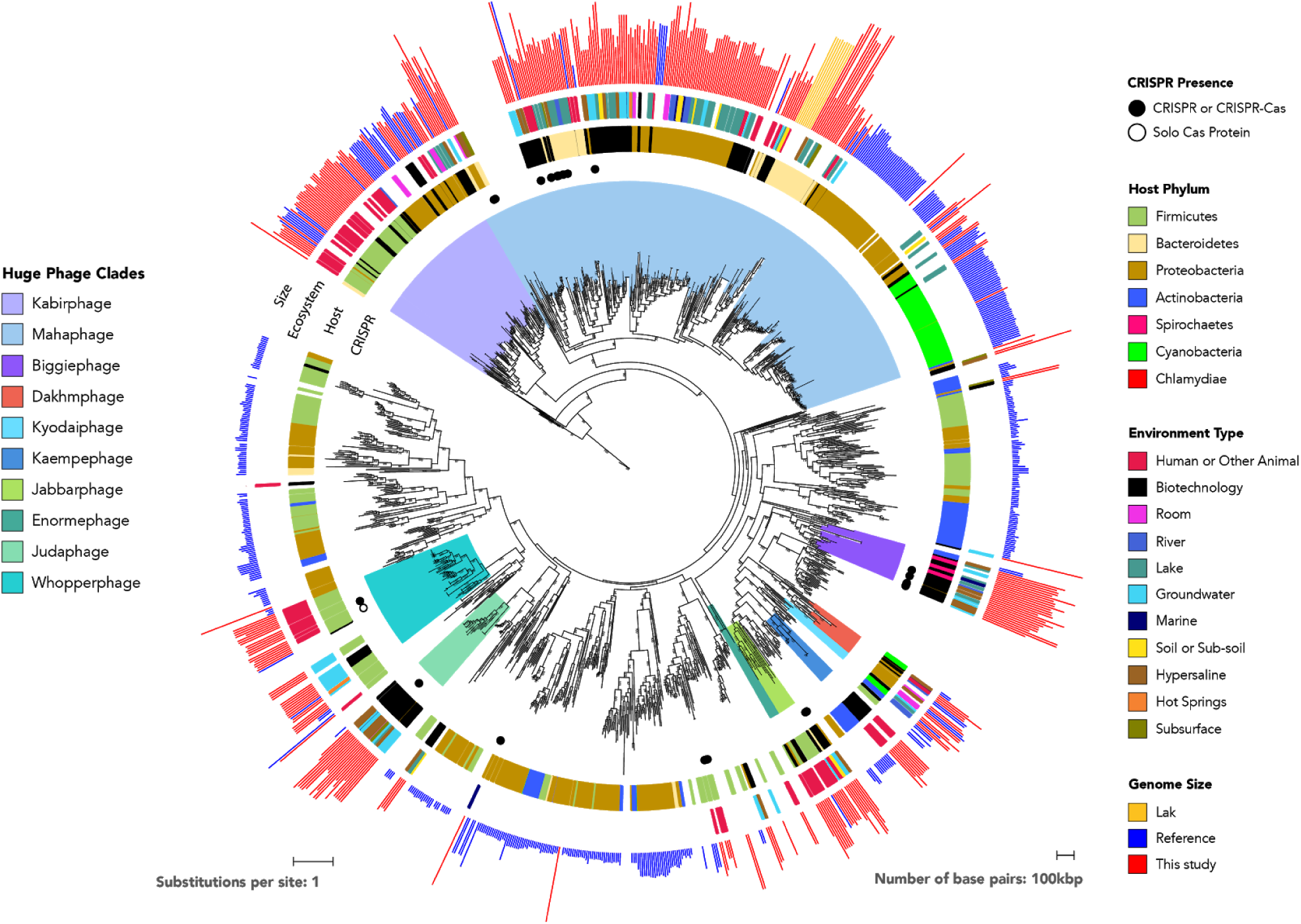
Phylogenetic reconstruction of huge phage evolutionary history using large terminase sequences from this study and similar matches from all RefSeq r92 proteins. The tree also includes large terminase sequences from complete and incomplete RefSeq phage, the Lak phage clade (Devoto *et al.*, 2019) and nonartifactual phage genomes that are >200 kbp from (Paez-Espino *et al.*, 2016). Huge phage clades identified in this study were independently corroborated with a phylogenetic reconstruction of major capsid genes (Fig S5A) and protein clustering (Fig S5B). The tree was rooted using 13 eukaryotic Herpesvirus terminases. The inner to outer rings display the presence of CRISPR-Cas from this study, host phylum, environmental sampling type, and genome size. Host phylum and genome were not included for RefSeq protein database matches because the sequence may be integrated prophage or part of organismal genome projects.

To determine the extent to which bacterial host phylogeny correlates with phage clades, we identified some phage hosts using CRISPR spacer targeting from bacteria in the same or related samples and phylogenies of normally host-associated genes that occur on phage (see below). We also tested the predictive value of bacterial taxonomic affiliations of the phage gene inventories (Methods) and found that in every case, CRISPR spacer targeting and phylogeny agreed with phylum-level taxonomic profiles. Consequently, we used the taxonomic profiles to predict the bacterial host phylum for many phage (**Table S3**). The results establish the importance of Firmicutes and Proteobacteria as hosts, and indicate the higher prevalence of Firmicutes huge phage in the human and animal gut compared to other environments (**Table 1**). Notably, the five genomes >634 kbp in length are all for phage predicted to replicate in Bacteroidetes, as do Lak phage (Devoto *et al.*, 2019), and all cluster within Mahaphage. Overall, phage grouped together phylogenetically are predicted to replicate in bacteria of the same phylum (**Figure 2**).

## Metabolism, transcription, translation

The phage genomes encode proteins predicted to localize to the bacterial membrane or cell surface. These may impact the susceptibility of the host to infection by other phage (**Table S4** and SI). We identified almost all previously reported categories of genes suggested to augment host metabolism during infection. Many phage have genes involved in *de novo* biosynthesis of purines and pyrimidines, and the interconversion of nucleic and ribonucleic acids and nucleotide phosphorylation states. These gene sets are intriguingly similar to those of bacteria with very small cells and putative symbiotic lifestyles (Castelle *et al.*, 2018) (**Table S4**).

Notably, many phage have genes whose predicted functions are in transcription and translation (**Table S5)**. Phage encode up to 64 tRNAs per genome, with sequences distinct from those of their hosts (**Table S6**). Generally, the number of tRNAs per genome increases with genome length (**Figure 1**). They have up to 16 tRNA synthetases per genome (**Table S6**), also distinct from but related to those of their hosts (**Figure S6** and SI). Phage may use these proteins to charge their own tRNA variants with host-derived amino acids. A subset of genomes have genes for tRNA modification and ligation of tRNAs cleaved by host defenses.

Many phage carry genes implicated in translation initiation, suggesting interception and redirection of host translation. These genes include initiation factors IF1 and IF3, as well as ribosomal proteins S1 and S21 (a phenomenon only recently reported in phage (Mizuno *et al.*, 2019); **Figure 3**). Both ribosomal proteins are important for translation initiation in bacteria (Van Duin and Wijnands, 1981; Farwell *et al.*, 1992; Sørensen *et al.*, 1998), making them likely useful for the hijacking of host ribosomes. Further analysis of S21 proteins revealed N-terminal extensions rich in basic and aromatic residues important for RNA binding. We predict that these phage ribosomal proteins substitute for host proteins (Mizuno *et al.*, 2019), and their extensions assist in competitive ribosome binding or preferential initiation of phage mRNAs.

**Figure 3:**
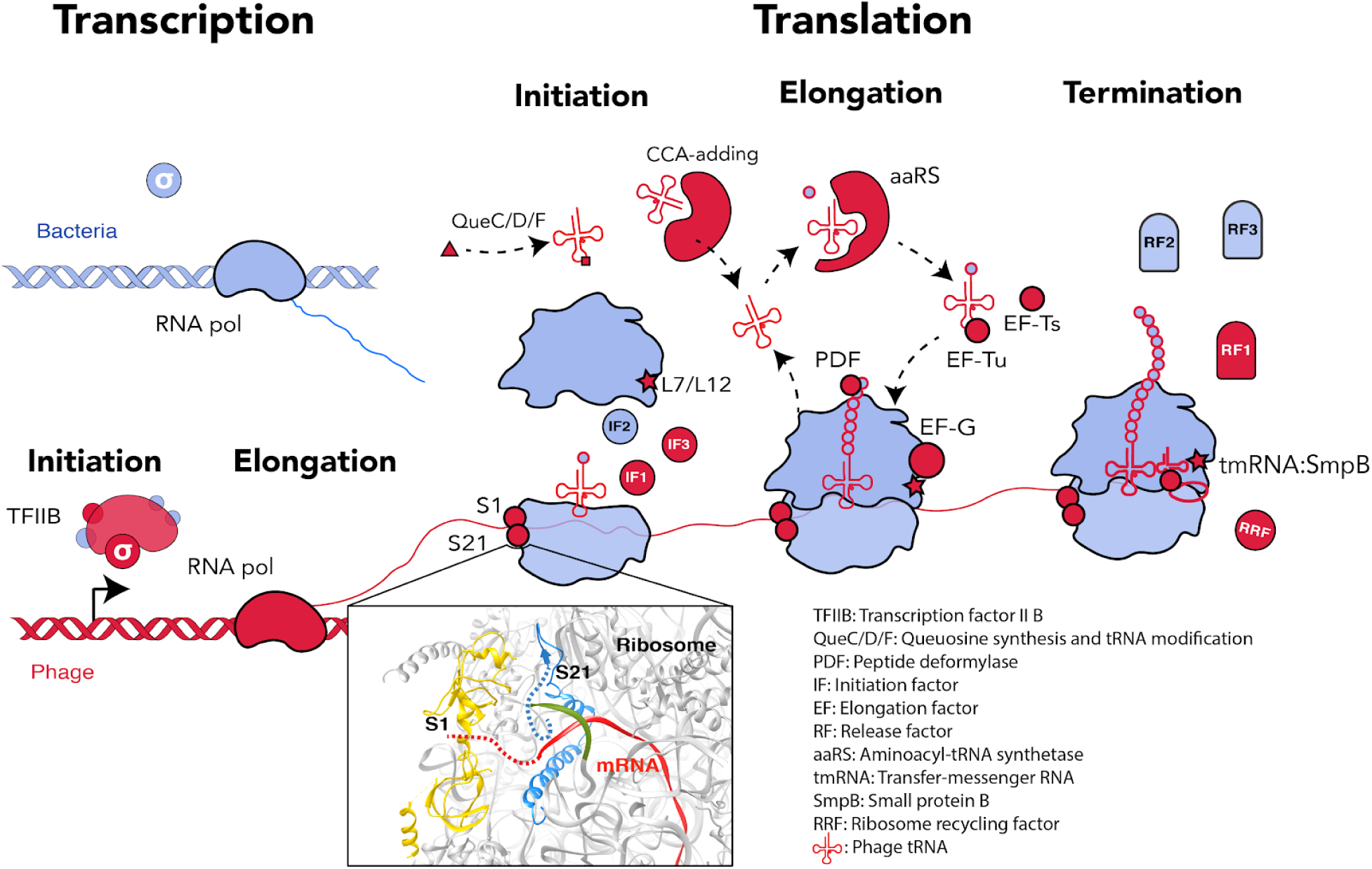
Model for how phageen-coded capacities could function to redirect the host’s translational system to produce phage proteins (bacterial components in blue, phage in red). No huge phage has all of these genes, but many have tRNAs and tRNA synthetases (see **Table S6**). Phage proteins with up to 6 ribosomal protein S1 domains occur in a few genomes. The S1 binds mRNA to bring it into the site on the ribosome where it is decoded (Subramanian, 1983). Phage ribosomal protein S21 might promote translation initiation of phage mRNAs, and many sequences have Nterminal extensions that may be involved in binding RNA (dashed blue line in ribosome insert, PDB: 6BU8 (Loveland and Korostelev, 2018), analyzed with UCSF Chimera (Pettersen *et al.*, 2004). Many other proteins of the translational apparatus are encoded by huge phage, belonging to all steps of the translation cycle.

Because protein S1 is often studied in the context of Shine Dalgarno (SD) sequence recognition by the ribosome (Farwell *et al.*, 1992; Sørensen *et al.*, 1998), we predicted the ribosomal binding sites for each phage genome (Methods). While most phage have canonical SD sequences, huge phage from this study that carry possible rpS1s rarely have identifiable SD sequences (**Figure S7** and **Table S7**). It is difficult to confirm “true” ribosomal S1 proteins due to the ubiquity of the S1 domain, but this correlation with non-canonical SD sequences suggests a role in translation initiation, either on or off the ribosome.

While assuming control of initiation may be the most logical step for phage redirection of host translation, efficiency of elongation and termination is necessary for robust infection and replication. Accordingly, we found many genes associated with the latter steps of translation in phage genomes. These include elongation factors G, Tu, and Ts, rpL7/12, and the processing enzyme peptide deformylase (PDF) (**Figure 3**), previously reported in phage genomes (Frank *et* al., 2013). We hypothesize that phage-encoded elongation factors act to maintain overall translation efficiency during infection, much like PDF’s prior predicted role in sustaining translation of necessary host photosynthetic proteins (Frank *et al.*, 2013). Translation termination factors are also represented in our huge phage genomes, including release factor 1, ribosome recycling factor, as well as tmRNAs and small protein B (SmpB), which rescue ribosomes stalled on damaged transcripts and trigger the degradation of aberrant proteins. These tmRNAs are also used by phage to sense the physiological state of host cells and can induce lysis when the number of stalled ribosomes in the host is high (Janssen and Hayes, 2012). Interestingly, some large putative plasmids also have analogous suites of translation relevant genes (**Table S4**).

About half of the phage genomes have one to fifty sequences >25 bp in length that fold into perfect hairpins. The palindromes (sequences with dyad symmetry) are almost exclusively intergenic and each is unique within a genome. Some, but not all, are predicted to be Rhoindependent terminators, thus provide clues regarding genes that function as independently regulated units (Methods). However, some palindromes are larger: up to 74 bp in length, with 34 genomes having examples ≥40 nt in length. These may have alternative or additional functions, such as modulating translation initiation and processivity.

## CRISPR-Cas mediated interactions

We identified most major types of CRISPR-Cas systems on phage, including Cas9-based Type II, the recently described Type V-I (Yan *et al.*, 2019), new variants of Type V-U systems (Shmakov *et al.*, 2017), and new subtypes of Type V-F (Harrington *et al.*, 2018). The Class II systems (types II and V) are reported in phage for the first time. Most phage effector nucleases (for interference) have conserved catalytic residues, implying that they are functional (**Figure S8**).

Unlike the well-described case of a phage with a CRISPR system (Seed *et al.*, 2013), almost all phage CRISPR systems lack spacer acquisition machinery (Cas1, Cas2, and Cas4) and many lack recognizable genes for interference (**Figure 4A**). For example, two related phage have a Type I-C variant system lacking Cas1 and Cas2 and have a helicase protein in lieu of Cas3 (**Figure S9**). They also harbor a second system containing a new candidate ∼750 aa Type V effector protein, tentatively named Cas12J (**Figure 4B**), that occurs proximal to CRISPR arrays.

**Figure 4:**
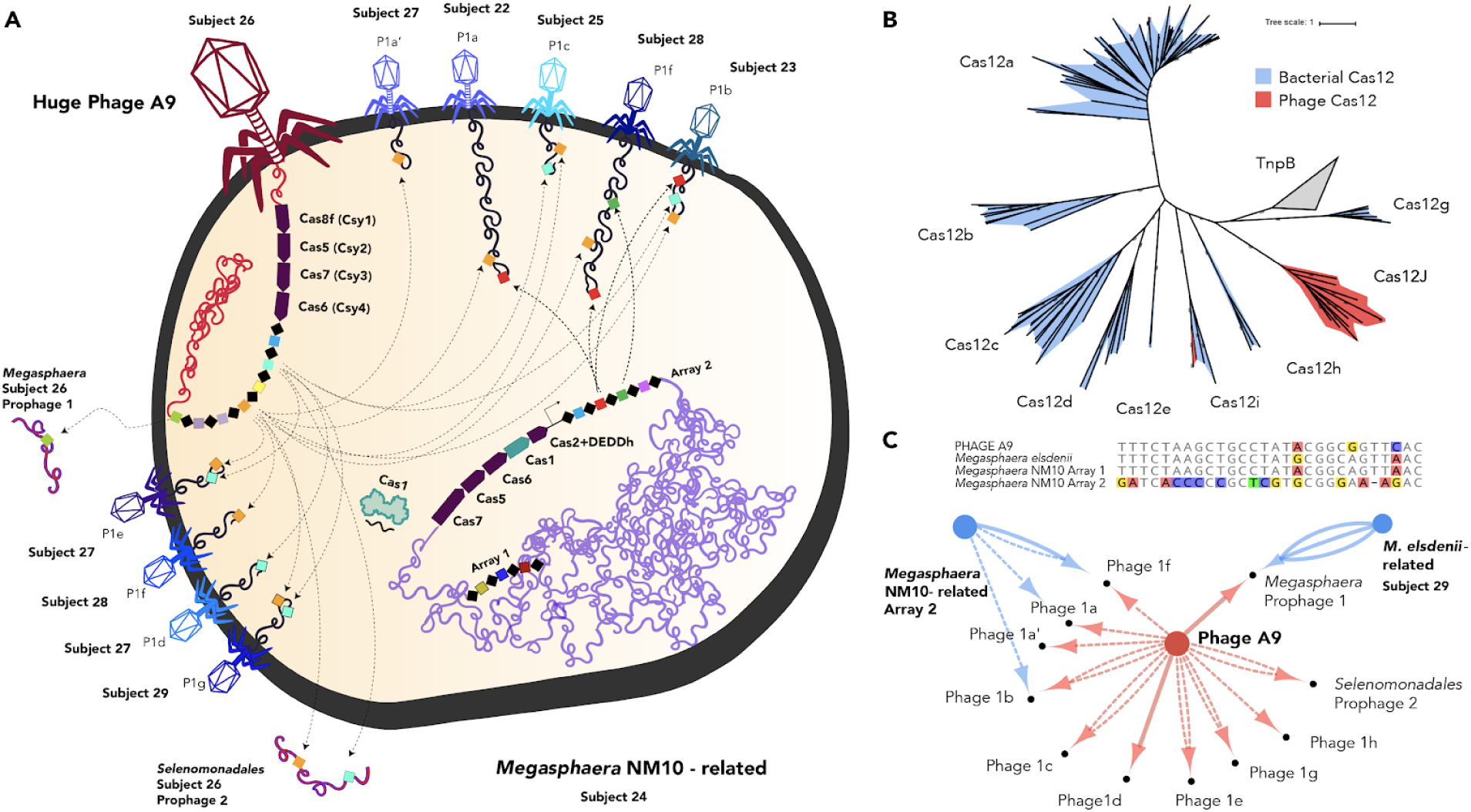
**A.** Cell diagram of bacterium-phage and phagephage interactions involving CRISPR targeting during superinfection. Arrows indicate CRISPR-Cas targeting of the prophage and phage genomes. Phage names indicate related groups delineated via whole genome alignment. We only included CRISPR interactions from samples of subjects of the same human cohort. **B.** Maximum likelihood phylogenetic tree of Cas12 subtypes a-i. Phageencoded Cas12i and Cas12J, the new effector, are outlined in red, with bacterialencoded proteins in blue. Bootstrap values above 90 are shown on the branches (circles). Cas14 and Type V-U trees are provided separately in the SI. **C.** Top panel shows the alignment of the consensus repeats from the A9 phage array and predicted host bacterial arrays. Bottom panel is an interaction network showing targeting of bacterial- (blue) and phage- (red) encoded CRISPR spacers. Number of edges indicate number of spacers from the array with targets to the smaller node. Solid edges denote spacer targets with no or 1 mismatch, and dashed edges denote 2-3 mismatches (to account for degradation in oldend phage spacers, diversity in different subjects, or phage mutation to avoid targeting).

In some cases, phage lacking genes for interference and spacer integration have similar CRISPR repeats as their hosts (**Figure 4C**), and thus may utilize Cas proteins synthesized by their host. Alternatively, systems lacking an effector nuclease may repress transcription of the target sequences without cleavage (Luo *et al.*, 2015; Stachler and Marchfelder, 2016), or spacer-repeat guide RNAs may act in an RNAi-like mechanism to silence host CRISPR systems or nucleic acid they can hybridize to. The phageencoded CRISPR arrays are often compact (3-23 repeats; median 6 per array; **Figure S11**). This range is substantially smaller than typically found in prokaryotic genomes (Toms and Barrangou, 2017). Some phage spacers target core structural and regulatory genes of other phage (**Figure 4C**). Thus, phage apparently augment their hosts’ immune arsenal to prevent infection by competing phage (**Table S8**).

Some phage-encoded CRISPR loci have spacers that target bacteria in the same sample or in a sample from the same study. We suppose that the targeted bacteria are the hosts for these phage, an inference supported by other host prediction analyses (**Table S3**). Some loci with bacterial chromosome-targeting spacers encode Cas proteins that could cleave the host chromosome, whereas others do not. Targeting of host genes could disable or alter their regulation, which may be advantageous during the phage infection cycle. Some phage CRISPR spacers target bacterial intergenic regions, possibly interfering with genome regulation by blocking promoters or silencing non-coding RNAs.

Interesting examples of CRISPR targeting of bacterial chromosomes involve transcription and translation genes. For instance, one phage targets a σ^70^ in its host’s genome and encodes its own σ^70^ transcription factor (see SI). Some huge phage genomes encode anti-sigma factors (AsiA), consistent with prior reports of σ^70^ hijacking by phage with AsiA (Brown and Hughes, 1995). In another example, a phage spacer targets the host Glycyl tRNA synthetase, but the Cas14 effector lacks one of the required catalytic residues for cleavage, suggesting a role in repression (as a ‘dCas14’), rather than in cleavage (discussed in SI).

Interestingly, we found no evidence of host-encoded spacers targeting any CRISPR bearing phage. However, phage CRISPR targeting of other phage that are also targeted by bacterial CRISPR (**Figure 4C**) suggested phage-host associations that were broadly confirmed by the phage taxonomic profile (**Table S3**).

Some large *Pseudomonas* phage encode Anti-CRISPRs (Acr) (Bondy-Denomy *et al.*, 2015; Pawluk *et al.*, 2016) and proteins that assemble a nucleus-like compartment segregating their replicating genomes from host defense and other bacterial systems (Chaikeeratisak, Nguyen, Khanna, *et al.*, 2017). We identified proteins encoded in huge phage genomes that cluster with AcrVA5, AcrVA2, and AcrIIA7 and may function as Acrs. Also, identified were tubulin-homologs (PhuZ) and proteins (SI) that create a proteinaceous phage “nucleus” (Chaikeeratisak, Nguyen, Egan, *et al.*, 2017). The phage nucleus was recently shown to protect the phage genome against host defense by physically blocking CRISPR-Cas degradation (Mendoza *et al.*, 2018).

## Conclusions

We show that phage with huge genomes are widespread across Earth’s ecosystems. We manually curated, circularized and finished 34 novel phage genomes, distinguishing them from prophage, providing accurate genome lengths and complete inventories of genes, including those encoded in complex repeat regions that break automated assemblies. Even closely related phage have diversified across habitats, presumably following their bacterial hosts. Host and phage migration could transfer genes relevant in medicine and agriculture (e.g., pathogenicity factors and antibiotic resistance, SI). Additional medical significance could involve direct or indirect activation of immune responses. For example, some phage directly stimulate IFNg via a TLR9-dependent pathway and exacerbate colitis (Gogokhia *et al.*, 2019). Huge phage may represent a reservoir of novel nucleic acid manipulation tools with applications in genome editing and might be harnessed to improve human and animal health. For instance, huge phage equipped with CRISPR-Cas systems might be tamed and used to modulate bacterial microbiome function or eliminate unwanted bacteria.

The huge phage define massive clades, suggesting that a gene inventory comparable in size to those of many symbiotic bacteria is a conserved strategy for phage survival. Overall, their genes appear to redirect the host’s protein production capacity to favor phage genes by first intercepting the earliest steps of translation and then ensuring efficient protein production thereafter. These inferences are aligned with findings for some eukaryotic viruses, which control every phase of protein synthesis (Jaafar and Kieft, 2019). Some acquired CRISPR-Cas systems with unusual compositions that may function to control host genes and eliminate competing phage.

More broadly, huge phage represent little known biology, the platforms for which are distinct from those of small phage and partially analogous to those of symbiotic bacteria, somewhat blurring the distinctions between life and non-life. Given phylogenetic evidence for large radiations of huge phage, we wonder if they are ancient and arose simultaneously with cells and other phage from a pre-life (protogenote) state (Woese, 1998) rather than appearing more recently via episodes of genome expansion.

## Acknowledgments

Funding for this project was provided by National Institutes of Health (NIH) under awards RAI092531A and R01G-M109454 and the Alfred P. Sloan Foundation under grant APSF-20121005 to JFB and MM; National Science Foundation grant (NSF) Sustainable Chemistry grant 1349278 to JFB, NSF Graduate Research Fellowships to BAS (DGE 1752814) and MRO (DGE 1106400); the Paul Allen Foundation Frontiers Group; Chan Zuckerberg Biohub; Innovative Genomics Institute. The Novo Nordisk Foundation (NNF16OC0021856) to P.M; NASA 13R-0043 and CA AES to KRA and KF; German Science Foundation postdoctoral scholarship to A.J.P (DFG PR 1603/1-1); Camille & Henry Dreyfus Postdoctoral Fellowship in Environmental Chemistry to C.H.; Watershed Function Scientific Focus Area funded by the U.S. Department of Energy, Office of Science, Office of Biological and Environmental Research (DE-AC02-05CH11231) to A.L., P.M.C, and A.T; NSF grants (GRT00048468 and 1342701) to KCW and MB and (CHE-1740549) to JHDC; the March of Dimes Prematurity Research Center at Stanford University School of Medicine, the Thomas C. and Joan M. Merigan Endowment at Stanford University to DAR; the National Research Foundation of South Africa (UID 64877) to STLH and NSF CZO funding. We acknowledge the scientists who generated public database sequences and thank Allison Sharrar, Jenny Tung, Elizabeth Archie, Frank Aarestrup and Ricardo Kruger who contributed data. We thank Patrick Pausch and Gavin Knott for helpful discussions.

## Author contributions

Analyses were conducted primarily by BAS, RS, L-XC, FW and JFB. Specifically, phylogenetic and gene inventory analyses were performed by BAS and RS, tRNA synthetase analysis was led by L-XC, ribosome analyses were led by FW and BAS, and genome analysis scripts were written by RS. Size distribution analysis was conducted by RS. Genome curation was performed by JFB, PM, L-XC, AD, and CS. BAS led the CRISPR-Cas analyses. BAS, RS, and JFB wrote the manuscript, with input from all authors. Datasets were generated by the authors. The study was conceived by JFB, with input from PM and AD.

---

**Figure S1:**
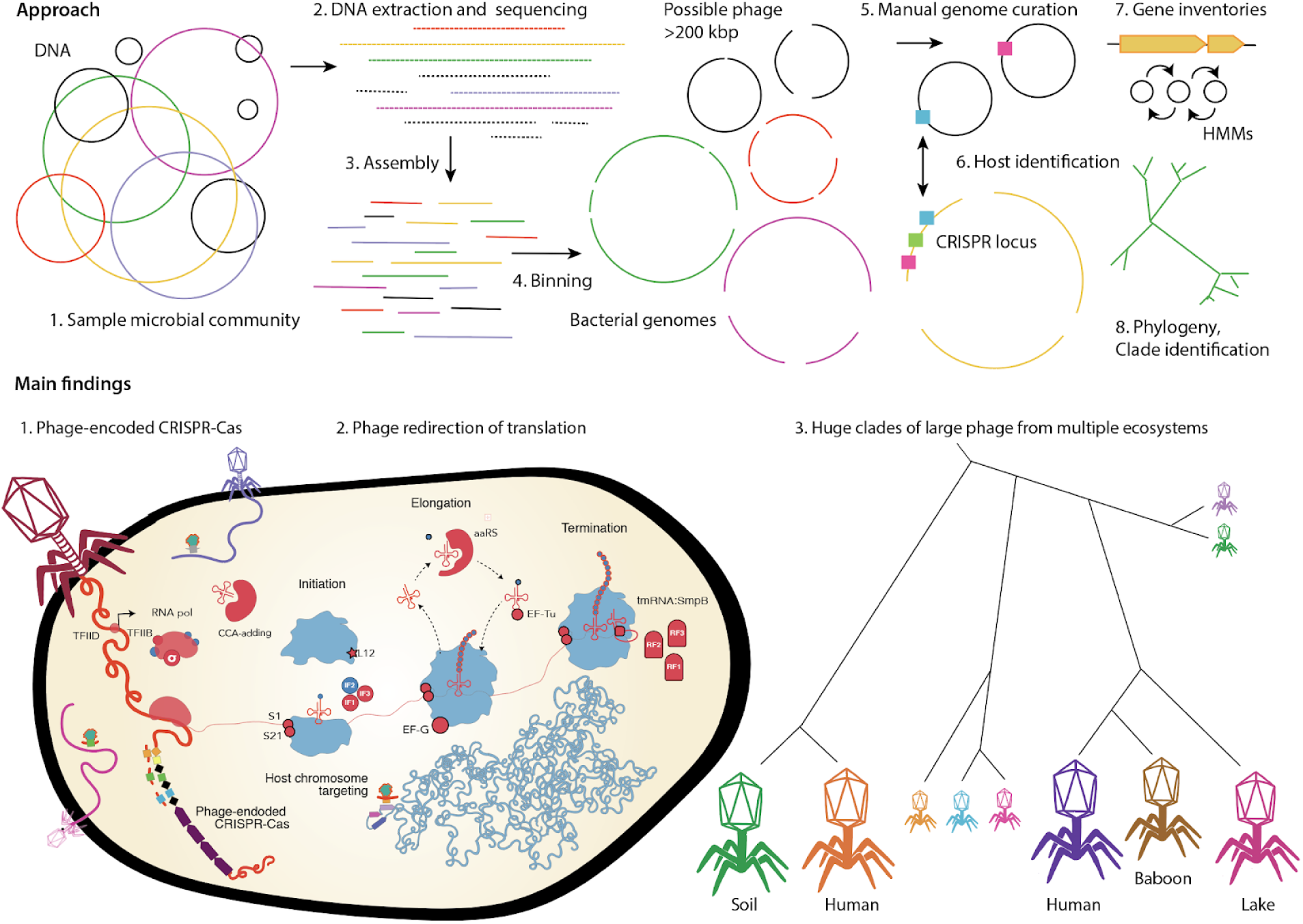
Graphical abstract describing the approach and main findings of this study.

## Methods

### Phage and plasmid genome identification

Datasets generated in the current study, those from prior research conducted by our team, the Tara Oceans microbiomes (Karsenti *et al.*, 2011), and the Global Oceans Virome (GOV) (Roux *et al.*, 2016) were searched for sequence assemblies that could have derived from phage with genomes of >200 kbp in length. Read assembly, gene prediction, and initial gene annotation followed standard methods reported previously (Joshi and Fass, 2011; Peng *et al.*, 2012; Edgar, 2015; Bushnell, 2016; Nurk *et al.*, 2017).

Phage candidates were initially found by retrieving sequences that were not assigned to a genome and had no clear taxonomic profile at the domain level. Taxonomic profiles were determined through a voting scheme, where there had to be a winner taxonomy >50% votes at each taxonomic rank based on UniProt and ggKbase (ggkbase.berkeley.edu) database protein annotations (Raveh-Sadka *et al.*, 2015). Phage were further narrowed down by identifying sequences with a high number of hypothetical protein annotations and/or the presence of phage structural genes, *e.g.*, terminase, capsid, tail, holin. All candidate phage sequences were checked throughout to distinguish putative prophage from phage. Prophage were identified based on a clear transition into genome with a high fraction of confident functional predictions, often associated with core metabolic functions, and much higher similarity to bacterial genomes. Plasmids were distinguished from phage based on matches to plasmid partitioning and conjugative transfer genes.

### Phage and plasmid genome manual curation

All scaffolds classified as phage or phage-like were tested for end overlaps indicative of circularization. Assembled sequences that could be perfectly circularized were considered potentially “complete”. Erroneous concatenated sequence assemblies were initially flagged by searching for direct repeats >5 kb using Vmatch (Kurtz, 2003). Potentially concatenated sequence assemblies were manually checked for multiple large repeating sequences using the dotplot and RepeatFinder features in Geneious v9. Sequences were corrected and removed from further analysis if the corrected length was < 200 kbp.

A subset of the phage sequences were selected for manual curation, with the goal of finishing (replacing all N’s at scaffolding gaps or local misassemblies by the correct nucleotide sequences and circularization). Curation generally followed methods described previously (Devoto *et al.*, 2019). In brief, reads from the appropriate dataset were mapped using Bowtie2 v2.3.4.1 (Langmead and Salzberg, 2012) to the *de novo* assembled sequences. Unplaced mate pairs of mapped reads were retained with shrinksam (github.com/bcthomas/shrinksam). Mappings were manually checked throughout to identify local misassemblies using Geneious v9. N-filled gaps or misassembly corrections made use of unplaced paired reads, in some cases using reads relocated from sites where they were mis-mapped. In such cases, mis-mappings were identified based on much larger than expected paired read distances, high polymorphism densities, backwards mapping of one read pair, or any combination of these. Similarly, ends were extended using unplaced or incorrectly placed paired reads until circularization could be established. In some cases, extended ends were used to recruit new scaffolds that were then added to the assembly. The accuracy of all extensions and local assembly changes were verified in a subsequent phase of read mapping. In many cases, assemblies were terminated or internally corrupted by the presence of repeated sequences. In these cases, blocks of repeated sequence as well as unique flanking sequence were identified. Reads were then manually relocated, respecting paired read placement rules and unique flanking sequences. After gap closure, circularization, and verification of accuracy throughout, end overlap was eliminated, genes were predicted, and the start moved to an intergenic region, in some cases suspected to be origin based on a combination of coverage trends and GC skew (Brown *et al.*, 2016). Finally, the sequences were checked to identify any repeated sequences that could have led to an incorrect path choice because the repeated regions were larger than the distance spanned by paired reads. This step also ruled out artifactual long phage sequences generated by end to end repeats of smaller phage, which occur in previously described datasets (see above and **Figure S3**).

### Structural and functional annotation

Following identification and curation of phage genomes, coding sequences (CDS) and Shine-Dalgarno ribosomal binding site (RBS) motifs were predicted with prodigal using genetic code 11 (-m -g 11 -p single). The resulting CDS were annotated as previously described by searching against UniProt, UniRef100, and KEGG (Wrighton *et al.*, 2014). Functional annotations were further assigned by searching proteins against PFAM r32 (Finn *et al.*, 2014), TIGRFAMS r15 (Haft *et al.*, 2013), and Virus Orthologous Groups r90 (vogdb.org). tRNAs were identified with tRNAscan-SE 2.0 (Lowe and Eddy, 1997) using the bacterial model. tmRNAs were assigned using ARAGORN v1.2.38 (Laslett and Canback, 2004) with the bacterial/plant genetic code.

Clustering of the CDS into families was achieved using a two-step procedure. A first protein clustering was done using the fast and sensitive protein sequence searching software MMseqs (Hauser *et al.*, 2016). An all-vs.-all sequences search was performed using evalue: 1 × 10^3^, sensitivity: 7.5 and coverage: 0.5. A sequence similarity network was built based on the pairwise similarities and the greedy set cover algorithm from MMseqs was performed to define protein subclusters. The resulting subclusters were defined as subfamilies. In order to test for distant homology, we grouped subfamilies into protein families using an HMM-HMM comparison. The proteins of each subfamily with at least two protein members were aligned using the result2msa parameter of mmseqs2, and from the multiple sequence alignments HMM profiles were built using the HHpred suite. The subfamilies were then compared to each other using HHblits (Remmert *et al.*, 2011) from the HHpred suite (with parameters -v 0 -p 50 -z 4 -Z 32000 -B 0 -b 0). For subfamilies with probability scores of ≥ 95% and coverage ≥ 0.50, a similarity score (probability × coverage) was used as weights of the input network in the final clustering using the Markov Clustering algorithm, with 2.0 as the inflation parameter. These clusters were defined as the protein families. Protein sequences were functionally annotated based on the accession of their best Hmmsearch match (version 3.1) (e-value cutoff 1 × 10^3^) against an HMM database constructed based on ortholog groups defined by the KEGG database (Kanehisa *et al.*, 2016) (downloaded on June 10, 2015). Domains were predicted using the same hmmsearch procedure against the PFAM r31 database (Finn *et al.*, 2014). The domain architecture of each protein sequence was predicted using the DAMA software (Bernardes *et al.*, 2016) (default parameters). SIGNALP (Petersen *et al.*, 2011) (version 4.1) (parameters: -f short -t gram+) and PSORT (version 3.0) (Peabody *et al.*, 2016) (parameters: -long -positive) were used to predict the putative cellular localization of the proteins. Prediction of transmembrane helices in proteins was performed using TMHMM (Krogh *et al.*, 2001) (version 2.0) (default parameters). Hairpins (palindromes, based on identical overlapping repeats in the forward and reverse directions) were identified using the Geneious Repeat Finder and located dataset-wide using Vmatch (Kurtz, 2003). Repeats >25 bp with 100% similarity were tabulated.

### Reference genomes for size comparisons

RefSeq r92 genomes were recovered by using the NCBI Virus portal and selecting only complete dsDNA genomes with bacterial hosts. Genomes from (Paez-Espino *et al.*, 2016) were downloaded from IMG/VR and only sequence assemblies labeled “circular” with predicted bacterial hosts were retained. Given the presence of sequences in IMG/VR that are based on erroneous concatenations (**Figure S3**), we only considered sequences from this source that are >200 kb, but a subset of these were removed as artifactual sequences.

### Alternative genetic codes

In cases where gene prediction using the standard bacterial code (code 11) resulted in seemingly anomalously low coding densities, potential alternative genetic codes were investigated. In addition to making a prediction using Fast and Accurate genetic Code Inference and Logo (FACIL) (Dutilh *et al.*, 2011), we identified genes with well defined functions (*e.g.*, polymerase, nuclease) and determined the stop codons terminating genes that were shorter than expected. We then re-predicted genes using GLIMMER3 v1.5 (Delcher *et al.*, 1999) and prodigal with TAG not interpreted as a stop codon. Other combinations of repurposed stop codons were evaluated, and candidate codes (e.g., code 6, with only one stop codon) were ruled out due to unlikely gene fusion predictions.

### Large terminase subunit and major capsid phylogenetic analysis

The large terminase subunit phylogenetic tree was constructed by recovering large terminases from the aforementioned protein clustering and annotation pipeline. CDS that matched with > 30 bitscore against PFAM, TIGRFAMS, and VOG were retained. Any CDS that had a hit to large terminase, regardless of bitscore, was searched using HHblits (Steinegger *et al.*) against the uniclust30_2018_08 database. The resulting alignment was then further searched against the PDB70 database. Remaining CDS that clustered in protein families with a large terminase HMM were also included after manual verification. Detected large terminases were manually verified using HHPred (Steinegger *et al.*) and jPred (Cole *et al.*, 2008). Large terminases from the > 200 kbp (Paez-Espino *et al.*, 2016) phage genomes and all >200 kbp complete dsDNA phage genomes from RefSeq r92 were also included by protein family clustering with the phage CDS from this study. The resulting terminases were clustered at 95% amino acid identity (AAI) to reduce redundancy using CD-HIT (Huang *et al.*, 2010). Smaller phage genomes were included by searching the resulting CDS set against the full Refseq protein database and retaining the top 10 best hits. Those hits that had no large terminase match against PFAM, TIGRFAMS, or VOG were removed from further consideration and the remaining set was clustered at 90% AAI. The final set of large terminase CDS that were >100 aa were aligned using MAFFT v7.407 (--localpair --maxiterate 1000) (Katoh and Standley, 2013), and poorly aligned sequences were removed and the resulting set was realigned. The phylogenetic tree was inferred using IQTREE v1.6.6 using automatic model selection (Nguyen *et al.*, 2015). The phylogenetic tree of major capsid protein genes was constructed by retrieving all major capsid proteins annotated by combining the PFAM annotations of protein families and direct annotations by PFAM, TIGRFAMS, and VOG. Reference major capsid gene sequences were collected using the same strategy and sources as for the large terminase subunit tree. The resulting set were further screened by searching against PFAM, TIGRFAMS, and VOG and removing matches that had no large terminase match regardless of bitscore. The final set of major capsid sequences were aligned with MAFFT(--localpair --maxiterate 1000) and the phylogenetic tree was constructed using IQTREE with automatic model selection and 1000 bootstrap replicates.

### Whole genome scale clustering

To identify phage genomes that were closely related at the whole genome level we compared sequences using whole genome alignments. The goal of this analysis was to further corroborate the identified phylogenetic clades and test for the presence of very similar phages in different habitats and environments. Genomes grouped together in the primary clusters from dRep v2 (Olm *et al.*, 2017) were evaluated for genome alignment using Mauve (Darling *et al.*, 2004) within Geneious v9.

### CRISPR-Cas Locus and target detection

Phage and host encoded CRISPR loci (repeats and spacers) were identified using a combination of MinCED (github.com/ctSkennerton/minced) and CRISPRDetect (Biswas *et al.*, 2016). A custom database of Cas genes was built by collecting Cas gene sequences from (Makarova *et al.*, 2015; Shmakov *et al.*, 2015; Burstein *et al.*, 2017; Smargon *et al.*, 2017; Harrington *et al.*, 2018; Yan *et al.*, 2018, 2019) and built with MAFFT (--localpair --maxiterate 1000) and hmmbuild. CDS from this study were searched against the HMM database using hmmsearch with e-value < 1 × 10^-5^. Matches were checked using a combination of hmmscan and BLAST searches against the NCBI nr database and manually verified by identifying co-located CRISPR arrays and Cas genes. Spacers extracted from between repeats of the CRISPR locus were compared to sequences assemblies from the same site using BLASTN-short (Altschul *et al.*, 1990) Matches with alignment length >24 bp and ≤1 mismatch were retained and targets were classified as bacterial, phage, or other. CRISPR arrays that had at least one ≤1 mismatch, were further searched for more spacer matches in the target sequence by finding more hits with ≤3 mismatches.

### Host identification

The phylum affiliations of bacterial hosts for phage and plasmid-like sequences were predicted by considering the UniProt taxonomic profiles of every CDS for each phage genome. The phylum level matches for each phage genome were summed and the phylum with the most hits was considered as the potential host phylum. However, only cases where this phylum that had 3x as many counts as the next most counted phylum were assigned as the tentative phage host phylum. Phage hosts were further assigned and verified using the aforementioned CRISPR targeting strategy with the phage and plasmid-like genomes as targets. CRISPR arrays were predicted on all sequence assemblies from the same site that each phage genome was reconstructed. Sequence assemblies containing spacers with a match of length >24 bp and ≤1 mismatch. In the case of phage, the match was used to infer a phage-host relationship. In all cases, the predicted host phylum based on taxonomic profiling and CRISPR targeting were in complete agreement. Similarly, the phyla of hosts were predicted based on phylogenetic analysis of phage genes also found in host genomes (e.g., involved in translation and nucleotide reactions). Inferences based on computed taxonomic profiles and phylogenetic trees were also in complete agreement.

### Phage encoded tRNA synthetase trees

Phylogenetic trees were constructed for phage encoded tRNA synthetase, ribosomal, and initiation factor protein sequences using a set of the closest reference sequences from NCBI and bacterial genomes from the current study. The tRNA synthetases were identified based on annotation of genes via the standard ggKbase pipeline (see above), and confirmed by HMMs with datasets from TIGRFAMs. For each type of tRNA synthetase, references were selected by comparing all the corresponding genes of this type against NCBI nr using DIAMOND v0.9.24 (Buchfink *et al.*, 2015), their top 100 hits were clustered by CD-HIT with 90% similarity threshold (Huang *et al.*, 2010). The phylogenetic tree of each tRNA synthetase was constructed using RAxML v8.0.26 (Stamatakis, 2014) with the PROTGAMMALG model.

## Data Availability

Reads are being deposited in the short read archive (if not already lodged there) and genome sequences will be available in NCBI shortly.

